# The Limitations of TabPFN for High-Dimensional RNA-seq Analysis

**DOI:** 10.1101/2025.08.15.670537

**Authors:** Summer Zhou, Vinayak Agarwal, Ashwin Gopinath, Timothy Kassis

## Abstract

Tabular Prior-Data Fitted Networks (TabPFN) demonstrate remarkable performance on small-to-medium tabular datasets through in-context learning, but struggle with high-dimensional genomic data such as RNA-seq with tens of thousands of features. We investigate multiple approaches to adapt TabPFN for transcriptomic analysis using two benchmark datasets: Age-ARCHS4, a regression dataset derived from the ARCHS4 dataset (57,873 samples, 10,000 genes), and an Inflammatory Bowel Disease (IBD) classification dataset encompassing Crohn’s Disease and Ulcerative Colitis samples (2,490 samples, 10,000 genes). Our experimental design proceeds in two phases: first evaluating existing optimization methods, then testing novel adaptations including (1) self-supervised embedding learning and (2) BulkFormer integration. We demonstrate that when constrained to equal training conditions (500 features, 10,000 samples), TabPFN outperforms classical baselines like random forest and XGBoost. However, when classical methods utilize full feature sets while TabPFN adaptations attempt to handle higher-dimensional data, all TabPFN variants consistently underperform the naive baseline. Our findings reveal fundamental limitations in current approaches to adapting TabPFN for genomic applications, showing that architectural modifications paradoxically degrade performance, while intelligent metadata-based subgrouping emerges as the most effective strategy for deploying TabPFN on biological data.

## 1 Introduction

Tabular Prior-Data Fitted Networks (TabPFN) represent a breakthrough in tabular machine learning, offering state-of-the-art performance through transformer-based in-context learning without requiring gradient updates for new datasets [Hollmann et al., 2023]. When operating within its design constraints of ≤500 features and ≤ 10,000 samples, TabPFN consistently outperforms classical machine learning baselines. However, our dataset contains 10,000 genes and nearly 60,000 samples, substantially exceeding these architectural limits. When TabPFN is restricted to 500 randomly selected genes while classical baselines can utilize all available genes and samples, TabPFN’s performance advantage disappears. This performance gap motivated our investigation into self-supervised learning and domain-specific architectural modifications to enable TabPFN to effectively process high-dimensional genomic data, though as we demonstrate, these adaptations paradoxically underperform the naive TabPFN baseline.

This limitation is especially problematic in computational biology, where datasets are high-dimensional, sample sizes are small, and the data exhibit complex biological structure—such as gene co-expression networks, metabolic pathways, and regulatory relationships—that cannot be adequately captured when restricted to TabPFN’s 500-gene input constraint.

We address these challenges through two complementary approaches: (1) **Method I** employs self-supervised learning to compress RNA-seq profiles into 500-dimensional embeddings that preserve biological signal while fitting TabPFN’s architectural constraints, and (2) **Method II** integrates BulkFormer, a recently proposed transformer architecture explicitly designed for bulk transcriptomic data [Kang et al., 2025], which captures biological structure through graph-based modeling of gene-gene interactions. We adapt BulkFormer embeddings for downstream use with TabPFN to explore whether biologically enriched representations can improve generalization.

Our contributions include:

- Novel combination of self-supervised objectives with TabPFN, including masked gene modeling, contrastive learning, and variational autoencoders
- An integration of BulkFormer [Kang et al., 2025], a domain-specific encoder that captures gene-gene interactions and biological priors in bulk RNA-seq data
- Comprehensive evaluation on age prediction (regression) and inflammatory bowel disease classification tasks
- Empirical analysis of TabPFN’s scalability.

## 2 Related Work

### 2.1 TabPFN Extensions

Several recent works have attempted to scale TabPFN beyond its original limitations. TabICL [Qu et al., 2025] introduces two-stage row/column attention mechanisms supporting up to 500k rows and 500 features for classification tasks. TabFlex [Zeng et al., 2025] employs linear attention to handle millions of rows and thousands of features, also limited to classification. TabPFN Unleashed [Liu and Ye, 2025] proposes supervised fine-tuning of encoders but provides limited implementation details.

From these approaches, we adapt TabICL and TabFlex to our tasks. Since TabICL [Qu et al., 2025] focuses on extending the number of samples, but our classification dataset does not exceed TabPFN’s limitation of 10k samples, this method is not particularly useful for our classification task alone. However, we experiment with an architecture that first uses TabICL to categorize our regression dataset (which does exceed TabPFN’s limitation) into three age groups, then applies three separate TabPFN models for the respective predicted groups. TabFlex [Zeng et al., 2025], on the other hand, can effectively handle more of our features, which we test in our experiments.

### 2.2 Traditional and Modern Genomic Machine Learning Approaches

Classical machine learning for high-dimensional genomics, where the number of features far exceeds the number of samples, typically follows a pipeline that begins with preprocessing to reduce technical noise, such as normalization and batch-effect correction [Johnson et al., 2006]. Next, informative genes are selected either through statistical testing with multiple-testing control (Benjamini–Hochberg FDR) [Benjamini and Hochberg, 1995] or through embedded feature selection methods like LASSO, which imposes sparsity during model fitting [Tibshirani, 1996]. Dimensionality reduction methods, including PCA for capturing major variation[Jolliffe, 2011] and t-SNE for visualizing complex structures [van der Maaten and Hinton, 2008], are often applied to further reduce feature space complexity. The processed features are then used to train standard predictive models, such as Support Vector Machines [Cortes and Vapnik, 1995] or Random Forests [Breiman, 2001]. While these approaches can perform well on bulk transcriptomic data when combined with careful cross-validation and data leakage prevention, they rely heavily on manual feature engineering and are limited in their ability to model intricate interactions between genes and samples. In recent years, modern genomic ML has begun to shift toward large-scale foundation models trained directly on massive transcriptomic datasets. For example, BulkFormer [Kang et al., 2025] is a 150-million parameter transformer-based model pre-trained on over 500,000 bulk RNA-seq profiles. Such models aim to learn rich, general-purpose biological representations that can be fine-tuned for diverse downstream tasks. In this work, we systematically investigate how domain-specific genomic foundation models like BulkFormer can be integrated with TabPFN to address the challenges of high-dimensional genomic prediction.

## 3 Methods

### 3.1 Method I: Self-Supervised Embedding Learning

#### 3.1.1 Self-Supervised Objectives

**Masked Gene Modeling (MGM)**. The masked gene modeling pipeline employs a two-stage approach for genomic age prediction, combining self-supervised representation learning with TabPFN regression. The architecture consists of a deep encoder-decoder network that reduces dimensionality from approximately 10,000 genes to 500 latent features. The encoder comprises four fully-connected layers with layer normalization, ReLU activation, and dropout for regularization, while the decoder mirrors this structure in reverse. During self-supervised pre-training, 15% of gene expression values are randomly masked, and the model learns to reconstruct these masked genes from the remaining expression profile, encouraging the learning of biologically meaningful gene co-expression patterns. The learned 500-dimensional representations are subsequently fed into a TabPFN regressor for age prediction. This objective encourages the model to internalize gene–gene dependencies and co-expression structure. Conceptually related masked-token pretraining for single-cell data has been explored in Geneformer (predicting masked *gene identity* rather than expression) [Theodoris et al., 2023].

**Contrastive Learning** We implement a truly self-supervised contrastive learning approach that learns representations without using age labels during pre-training. The architecture employs a shared encoder with progressive compression using ReLU activation, batch normalization, and dropout, with *l*_2_-normalized output embeddings. To create positive pairs for contrastive learning, we apply three data augmentation strategies to each sample: (1) Gaussian noise injection with *σ* = 0.1, (2) random gene dropout with probability *p* = 0.1, and (3) expression scaling with factors uniformly sampled from [0.8, 1.2]. Each sample generates two augmented views that serve as positive pairs, while other samples in the batch constitute negative pairs. The model is trained using the InfoNCE loss function:

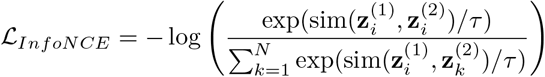

where **z***i*^(1)^ and 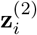 are *l*_2_-normalized embeddings of two augmented views of sample *i*, sim(·, *·*) denotes cosine similarity, *τ* = 0.1 is the temperature parameter, and *N* is the batch size. This approach encourages the model to learn invariant representations across biologically plausible data transformations while maintaining discriminative power for downstream age prediction.

**Variational Autoencoder** First introduced by Kingma and Welling [2013], the Variational Autoencoder (VAE) is a probabilistic generative model that learns a continuous, structured latent space. We implement a VAE architecture for probabilistic dimensionality reduction of gene expression data. The encoder follows a progressive compression structure with ReLU activation, batch normalization, and dropout, culminating in two separate linear layers that output the mean ***µ*** and log-variance ***σ***^2^ parameters of a 500-dimensional latent distribution. The decoder mirrors this architecture in reverse to reconstruct the original gene expression profile. The model is trained using the standard VAE objective:

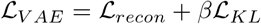

where 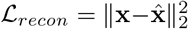 is the MSE reconstruction loss and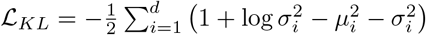 is the KL divergence regularization term. We use *β* = 0.1 to balance reconstruction fidelity with regularization. During inference, we sample from the learned latent distribution **z** *∼ N* (***µ***, diag(***σ***^2^)) to obtain 500-dimensional probabilistic representations. The VAE is optimized using AdamW with learning rate 5 *×* 10^*−*4^ and early stopping (patience = 15 epochs). This probabilistic approach captures uncertainty in the learned representations while encouraging a smooth, well-structured latent space suitable for downstream age prediction tasks.

**Deterministic Autoencoder** Originating from the work of Rumelhart et al. [1986] on using backpropagation to learn efficient data representations, we employ a deep autoencoder architecture for self-supervised dimensionality reduction of gene expression data. The encoder consists of four fully-connected layers with progressive compression, where each layer is followed by ReLU activation, batch normalization, and dropout (0.2) for regularization. The decoder mirrors this architecture in reverse to reconstruct the original gene expression profile. The model is trained using mean squared error (MSE) reconstruction loss:

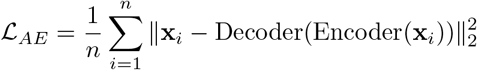

The autoencoder learns to compress high-dimensional gene expression data into a 500-dimensional latent representation that preserves the most informative biological patterns necessary for accurate reconstruction. We use the AdamW optimizer and early stopping with patience of 15 epochs. This self-supervised pretraining encourages the encoder to capture essential gene co-expression relationships without requiring age labels, producing biologically meaningful representations for downstream age prediction.

### 3.2 Method II: BulkFormer Integration

We evaluated BulkFormer [Kang et al., 2025], a 150-million parameter foundation model pre-trained on over 500,000 bulk RNA-seq profiles. BulkFormer employs a hybrid encoder architecture specifically designed for bulk transcriptomics that combines two complementary components: (1) a graph convolutional network (GCN) that captures explicit gene-gene interactions using pre-computed protein interaction networks derived from GTEx data, and (2) Performer modules that model global expression dependencies through efficient attention mechanisms with linear complexity. The model processes approximately 20,000 protein-coding genes simultaneously through three stacked GBFormer blocks, each containing graph convolutions followed by four Performer layers with 8 attention heads. Input gene expressions are embedded using three complementary representations: positional expression embeddings via RoPE-style encoding, pre-trained ESM2 protein embeddings, and direct expression-level encodings, which are summed and projected to a 640-dimensional hidden space.

Our integration pipeline leverages BulkFormer’s intermediate representations for downstream age prediction through a carefully designed preprocessing and dimensionality reduction strategy. Since BulkFormer’s pre-trained parameters depend on a fixed gene vocabulary and positional encoding established during its original training on 500,000+ bulk RNA-seq profiles, we align input gene expression data to match BulkFormer’s expected 20,010-gene sequence using the provided gene information file containing ENSEMBL gene identifiers, with missing genes filled using low expression values (−10). Gene expression data is normalized to log-transformed TPM values using gene length corrections and fed to the pre-trained BulkFormer model. We extract 640-dimensional contextualized embeddings from the second encoder layer, which captures biologically meaningful gene representations before the final reconstruction head. These gene-level embeddings are then aggregated to sample-level representations using mean pooling over approximately 4,000 high-variance genes identified during BulkFormer pre-training, resulting in 640-dimensional sample embeddings. To accommodate TabPFN’s 500-feature limit, we apply variance-based feature selection to identify the 500 most variable dimensions across the sample embeddings. The resulting 500-dimensional BulkFormer-derived features serve as input to TabPFN for age prediction, replacing traditional variance-based gene selection with foundation model representations that theoretically capture complex gene regulatory relationships and expression dependencies learned from large-scale transcriptomic data.

### 3.3 Method III: Existing TabPFN Optimization Approaches

Beyond the naive TabPFN baseline, we investigate several optimization strategies to enhance TabPFN’s performance on genomic data while respecting its architectural constraints of 500 features and 10,000 samples maximum.

#### TabPFN Ensembled

We implement an ensemble approach by training multiple TabPFN models on different subsets of the training data and averaging their predictions. Our ensemble consists of 15 independent TabPFN regressors, each trained on exactly 10,000 samples randomly sampled from the training pool to respect TabPFN’s sample limit. Each model uses a different random seed (corresponding to the run index) to ensure diverse sampling and model initialization. The training process maintains a fixed test set of 6,000 samples throughout all runs to enable consistent evaluation. For each individual model *k*, we train on samples {*X*_*k*_, *y*_*k*_*}* where |*X*_*k*_| = 10, 000, and the final ensemble prediction is computed as:

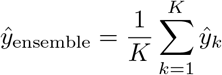

where *K* = 15 models and *ŷ*_*k*_ represents the prediction from the *k*-th TabPFN regressor. We evaluate cumulative ensemble performance after each model is added, allowing us to track improvement as ensemble size increases.

#### TabPFN Simple Average

For regression tasks, our majority vote approach adapts the ensemble methodology by computing mean predictions across multiple TabPFN models, which serves as the continuous analog to discrete majority voting. The implementation follows the same multi-model training procedure as the ensemble method, where 15 TabPFN regressors are trained on different random subsets of 10,000 samples each. The final prediction uses arithmetic mean aggregation of individual model outputs, effectively implementing a “soft” majority vote for continuous age predictions. This approach aims to reduce prediction variance and improve generalization through model diversity while maintaining the same computational framework as standard ensembling.

#### TabPFN Hierarchy

Inspired by TabICL [Qu et al., 2025]’s hierarchical approach, we implement a two-stage prediction system to handle the large sample size in our age dataset while respecting TabPFN’s constraints. The method first trains a TabPFN classifier to categorize samples into three age groups: young (1-35 years), middle (36-70 years), and old (71-114 years), using sophisticated feature selection that combines variance-based selection (30%), age correlation (40%), and mutual information (30%). After age group classification, we deploy three separate TabPFN regressors, each specialized for predictions within its respective age range. The hierarchical prediction process follows:

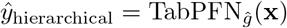

where *ĝ* = arg max_*g*_ TabPFN_classifier_(**x**) is the predicted age group and TabPFN_*ĝ*_ is the group-specific regressor trained only on samples within that age range. This approach allows the model to capture age-specific gene expression patterns while maintaining TabPFN’s sample size limitations, with each group-specific regressor trained on a subset that fits within the 10,000-sample constraint.

### 3.4 Datasets

- **Age-ARCHS4 (Regression)**: Derived from the ARCHS4 dataset, containing 57,873 bulk RNA-seq samples with 10,000 genes for chronological age prediction across diverse tissue types and conditions.
- **IBD Classification**: Inflammatory Bowel Disease dataset with 2,490 intestinal biopsy RNA-seq samples (10,000 genes) labeled as Ulcerative Colitis (UC), Crohn’s Disease (CD), or Control, representing a three-class classification task for gastrointestinal tract pathology.

### 3.5 Evaluation Protocol

We employ 5-fold stratified cross-validation with tissue type and age group stratification. Performance metrics include:

- **Regression**: Mean Absolute Error (MAE), *R*^2^
- **Classification**: Balanced Accuracy, AUROC, F1-score

### 3.6 Baselines

We compare against naive TabPFN, classical ML methods (Linear regression, Random Forest, XGBoost), TabPFN ensemble approaches (simple average, ensembling), and recent state-of-the-art TabPFN extensions (TabICL, TabFlex) specifically designed to overcome scalability limitations.

## 4 Results

### 4.1 Phase 1: Equal-Constraint Evaluation

We first evaluate TabPFN within its intended design constraints, comparing it against classical ML methods under identical training conditions (500 features selected by highest variance, 10,000 samples maximum). Figure 2 shows the predicted vs. actual age fit for TabPFN, while the Figure 3 summarizes model performance. Under these constraints, TabPFN achieves *R*^2^ = 0.606, outperforming Random Forest (*R*^2^ = 0.541) and XGBoost (*R*^2^ = 0.528), confirming its effectiveness in the regime it was designed for. This motivates further investigation into scaling strategies for larger genomic datasets.

**Figure 1.**
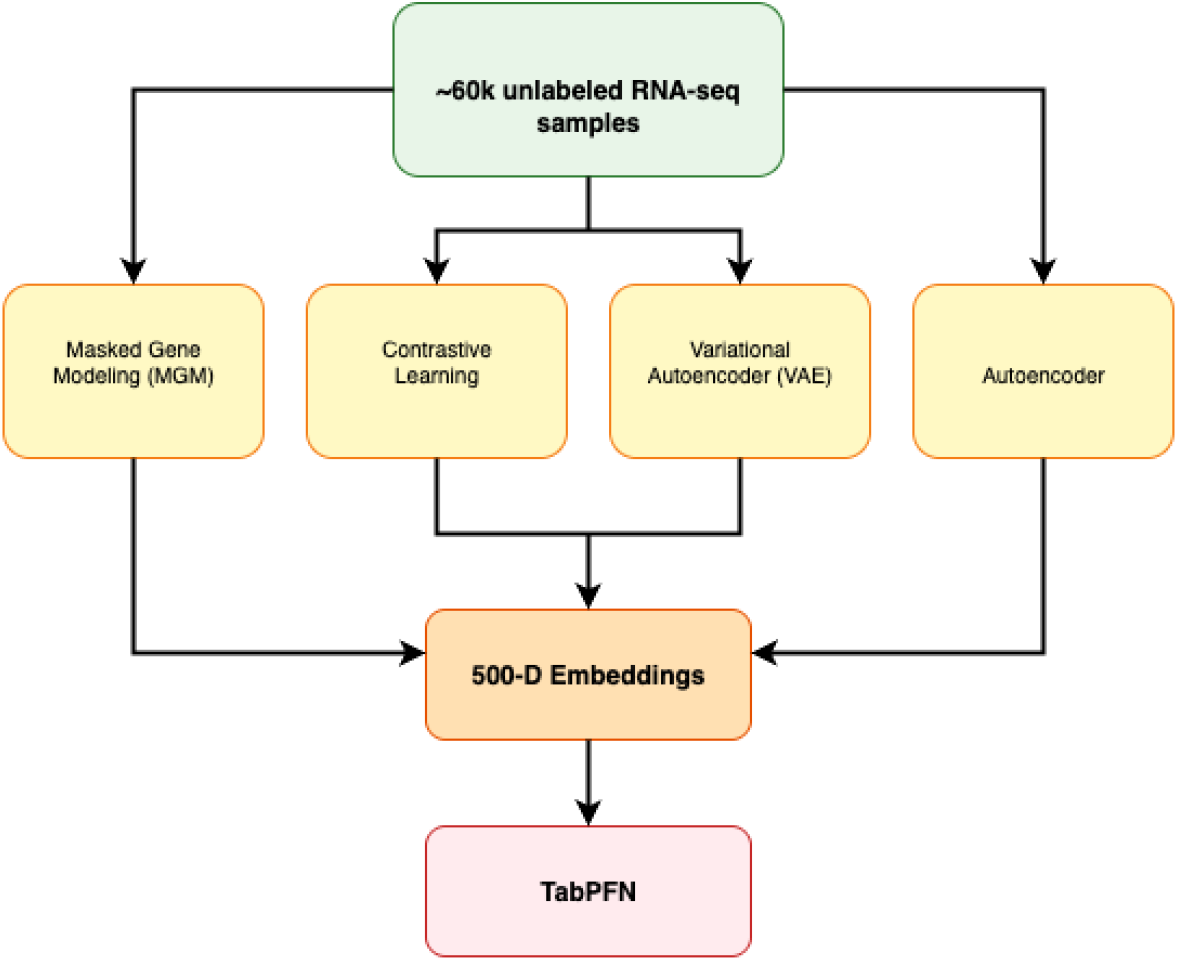
Method I Overview: Self-supervised embedding learning pipeline that compresses RNA-seq profiles into 500-dimensional representations for TabPFN integration.

**Figure 2.**
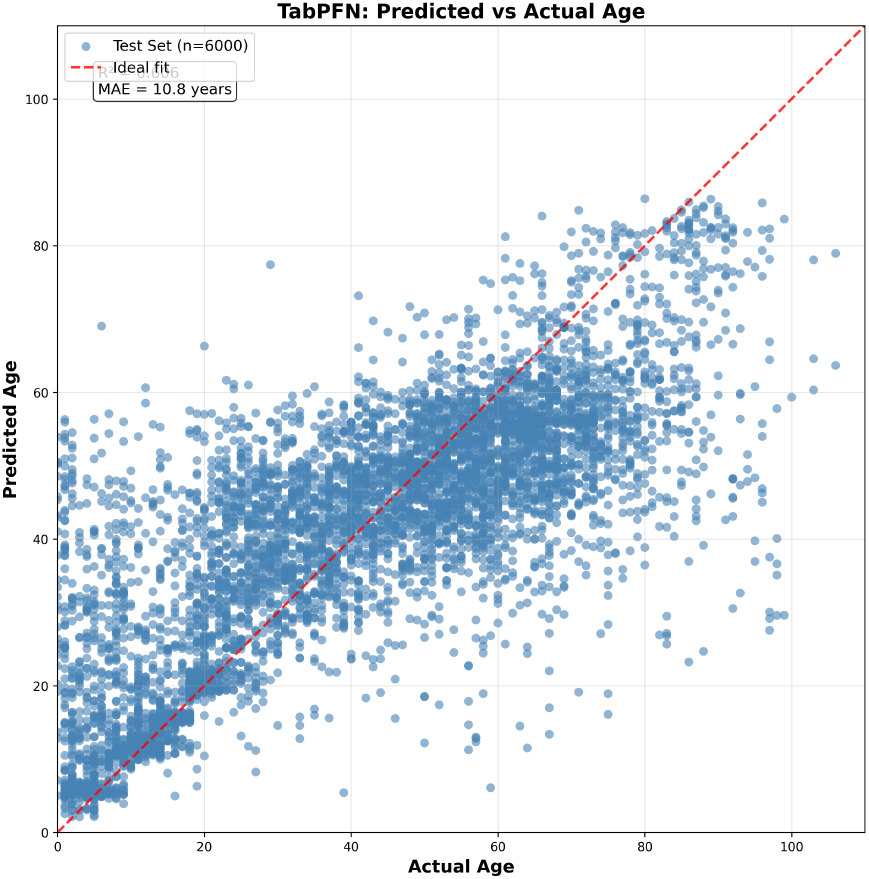
TabPFN predicted vs. actual age

**Figure 3.**
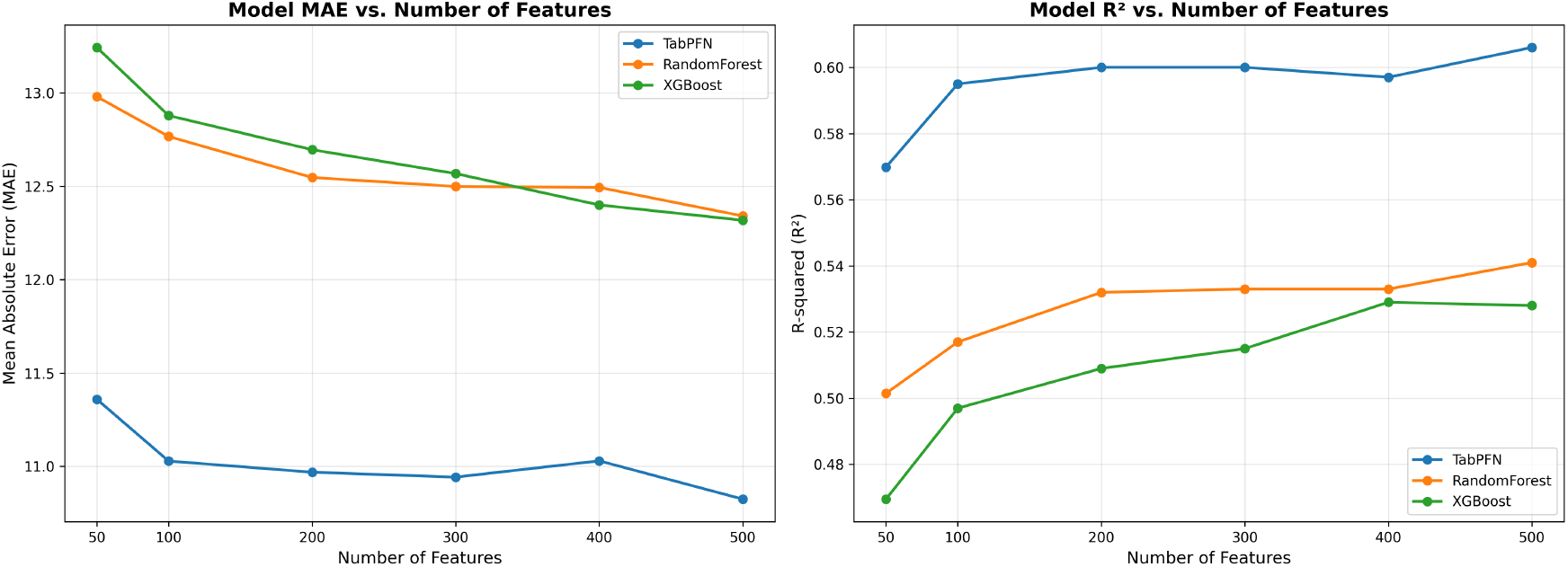
TabPFN performance vs. number of features. The steepest improvement occurs from 0 to 100 features, after which gains are incremental.

We also analyze the effect of feature count on TabPFN performance. As shown in Figure 3, the largest performance gain occurs when increasing from 0 to 100 features, with incremental returns beyond this point.

### 4.2 Phase 2: Adaptation Evaluation Results

We then evaluate various adaptation strategies as classical methods utilize full feature sets, testing: (1) existing optimization methods (ensemble approaches), (2) novel architectural adaptations (self-supervised embedding learning, BulkFormer integration), and (3) metadata-based subgrouping approaches.

#### 4.2.1 Age Prediction (Regression)

Table 2 and Figure 4 summarize age prediction performance when classical methods utilize full feature sets while various TabPFN adaptations attempt to handle high-dimensional data. Contrary to expectations, all TabPFN adaptations consistently underperformed the naive baseline.

**Table 1:** Model comparison results (500 features).

| Method | $R^2$ | MAE (years) |
| --- | --- | --- |
| *TabPFN* |  |  |
| TabPFN (500 features) | 0.606 | 10.824 |
| *Classical ML Baselines* |  |  |
| Random Forest | 0.541 | 12.341 |
| XGBoost | 0.528 | 12.318 |

**Table 2:**
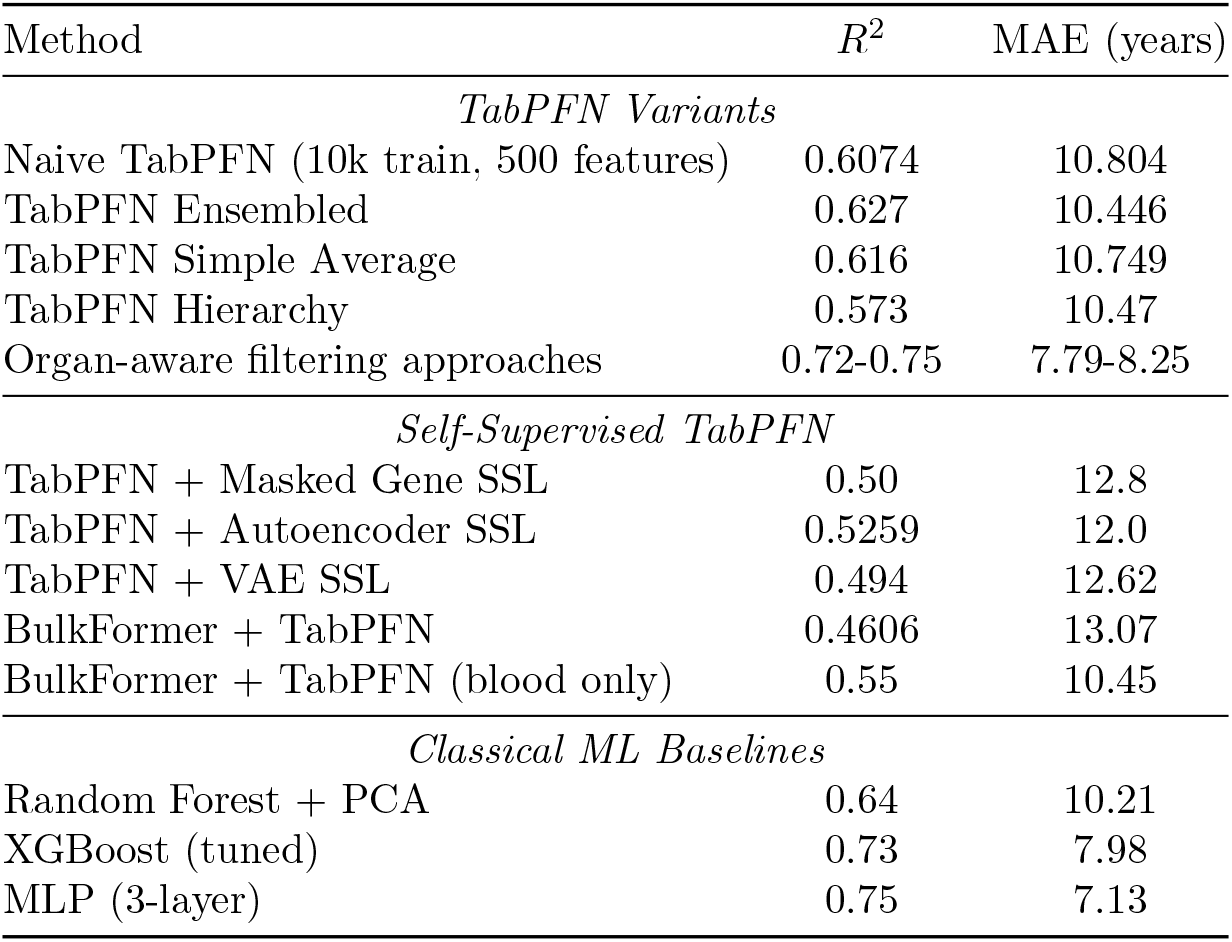
Age Prediction Results (Regression)

| Method | $R^2$ | MAE (years) |
| --- | --- | --- |
| *TabPFN Variants* |  |  |
| Naive TabPFN (10k train, 500 features) | 0.6074 | 10.804 |
| TabPFN Ensembled | 0.627 | 10.446 |
| TabPFN Simple Average | 0.616 | 10.749 |
| TabPFN Hierarchy | 0.573 | 10.47 |
| Organ-aware filtering approaches | 0.72-0.75 | 7.79-8.25 |
| *Self-Supervised TabPFN* |  |  |
| TabPFN + Masked Gene SSL | 0.50 | 12.8 |
| TabPFN + Autoencoder SSL | 0.5259 | 12.0 |
| TabPFN + VAE SSL | 0.494 | 12.62 |
| BulkFormer + TabPFN | 0.4606 | 13.07 |
| BulkFormer + TabPFN (blood only) | 0.55 | 10.45 |
| *Classical ML Baselines* |  |  |
| Random Forest + PCA | 0.64 | 10.21 |
| XGBoost (tuned) | 0.73 | 7.98 |
| MLP (3-layer) | 0.75 | 7.13 |

**Figure 4.**
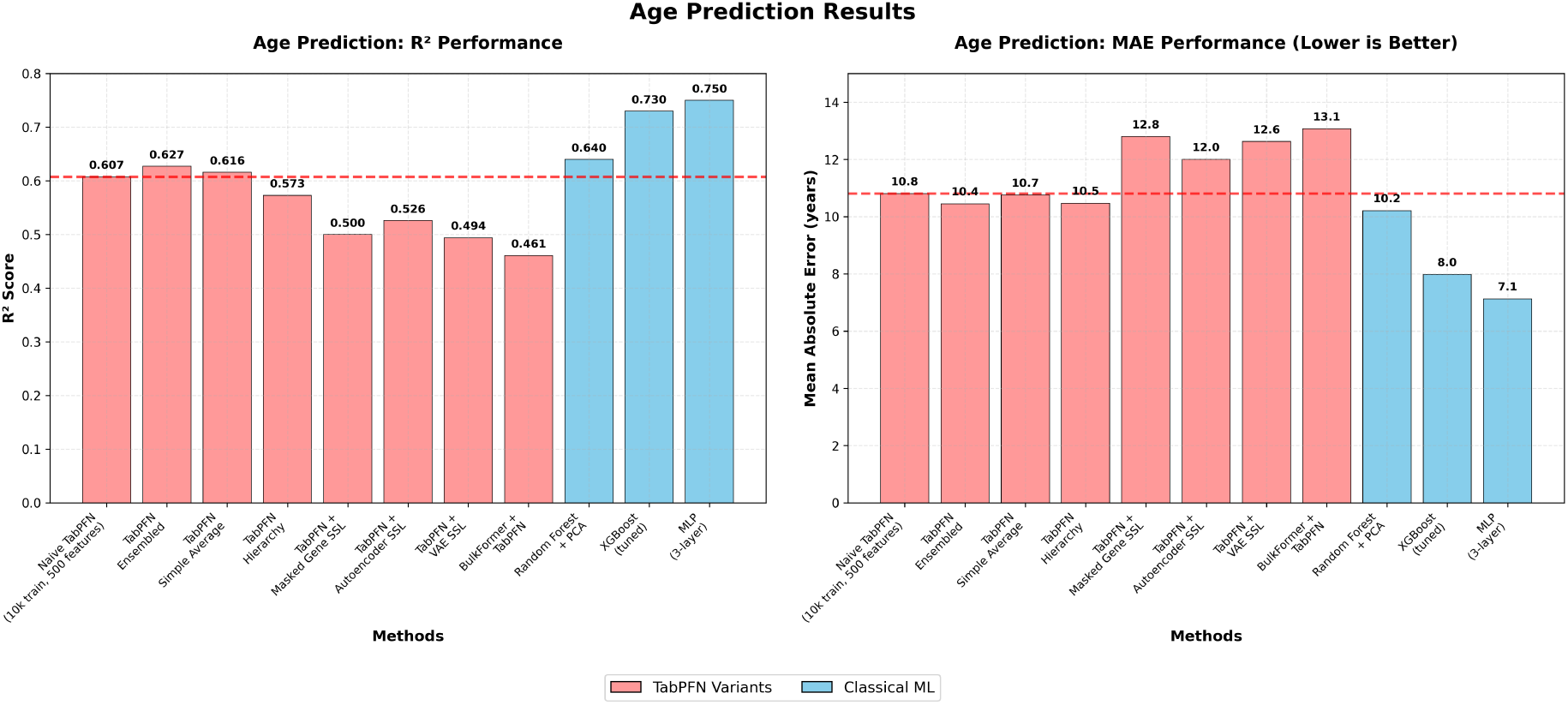
Age Regression Performance: MAE and *R*^2^ across all models. Left panel shows *R*^2^ scores, right panel shows Mean Absolute Error (lower is better). TabPFN variants (red bars) consistently underperform, while classical ML methods achieve superior results.

Key findings:

- **Most TabPFN adaptations underperformed naive TabPFN** (*R*^2^ = 0.607): hierarchy method achieved *R*^2^ = 0.573, while self-supervised variants showed particularly poor results (*R*^2^ = 0.494-0.5259)
- **BulkFormer integration reduced performance** (*R*^2^ = 0.4606), demonstrating poor compatibility between domain-specific architectures and TabPFN
- **Classical ML methods substantially outperformed** all TabPFN variants, with 3-layer MLP achieving *R*^2^ = 0.75 and MAE = 7.13 years—a 23% improvement over naive TabPFN
- Ensemble approaches (majority voting, ensembling) provided only marginal improvements (*R*^2^ = 0.616-0.627 vs 0.607)

#### 4.2.2 IBD Classification

Table 3 and Figure 5 present IBD classification results, showing similar patterns of TabPFN adaptation failure. Recent TabPFN extensions failed on genomic data: TabICL achieved only 57.8% balanced accuracy and TabFlex 51.5% on IBD classification, both substantially worse than naive TabPFN’s 63.7%.

**Table 3:** IBD Classification Results.

| Method | Balanced Accuracy | AUROC | F1-Score |
| --- | --- | --- | --- |
| *TabPFN Variants & Extensions* |  |  |  |
| Naive TabPFN (10k samples, 500 features) | 0.6367 | 0.8733 | 0.6385 |
| TabICL | 0.5781 | N/A | 0.7636 |
| TabFlex | 0.5154 | N/A | 0.6955 |
| *Self-Supervised TabPFN* |  |  |  |
| BulkFormer + TabPFN | 0.5442 | 0.77 | 0.5852 |
| Naive TabPFN (3 categories) | 0.70 | 0.86 | 0.718 |
| *Classical ML Baselines* |  |  |  |
| Random Forest + PCA | 0.50 | 0.80 | 0.52 |
| Random Forest + Filtering | 0.74 | 0.89 | 0.74 |

**Figure 5.**
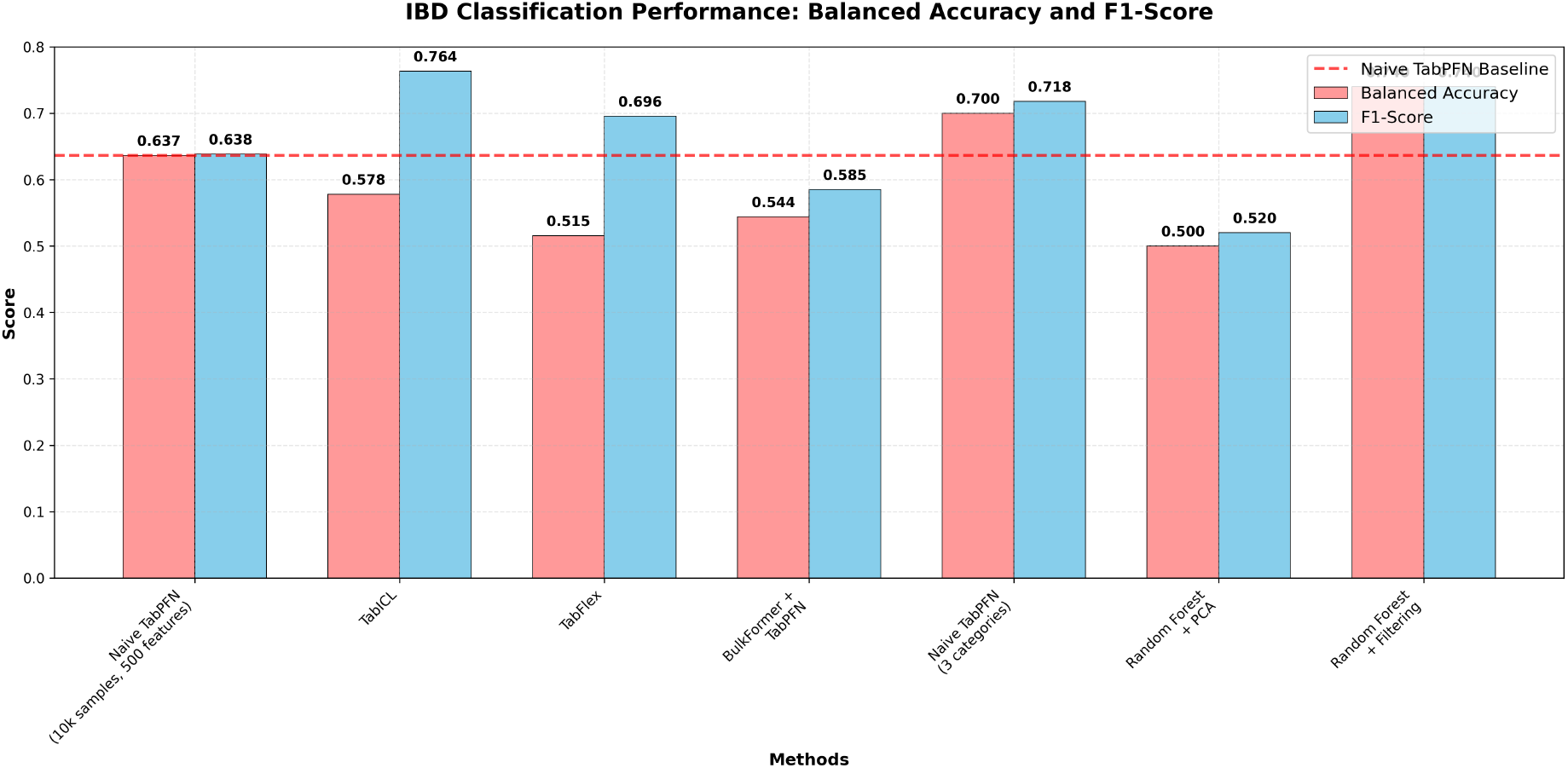
IBD Classification Performance: Balanced Accuracy and F1-Score across all models. TabPFN extensions (TabICL, TabFlex) and adaptations significantly underperform naive TabPFN baseline (dashed line), while classical methods achieve optimal results.

**Figure 6.**
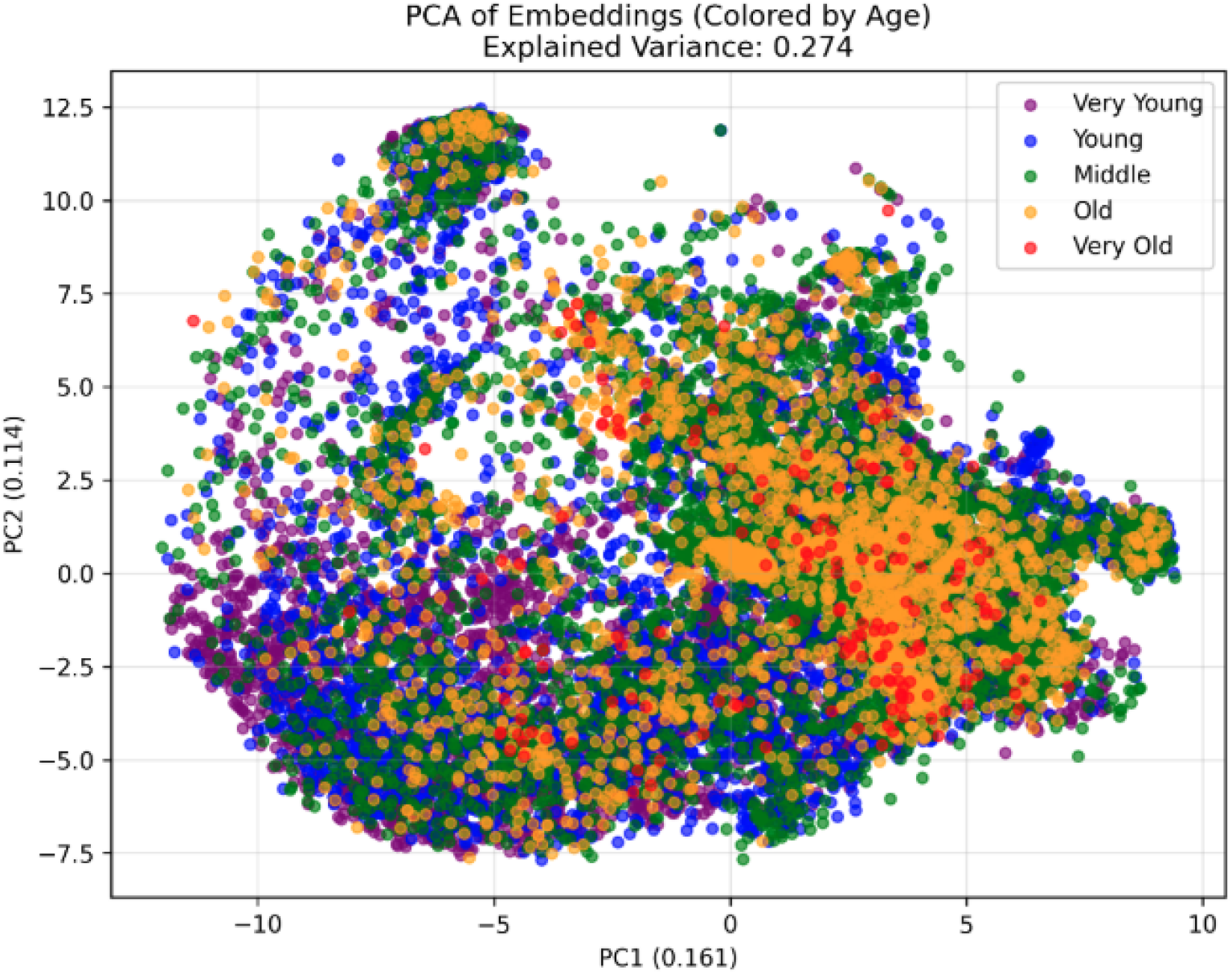
BulkFormer+TabPFN integration analysis. The age groups represented by BulkFormer are mostly overlapped, showing this is not a great way to dimensionally reduce on this dataset.

IBD classification results mirror the age prediction findings: TabPFN extensions performed worse (TabICL: 0.5781, TabFlex: 0.5154), and BulkFormer integration achieved only 0.5442 balanced accuracy. Notably, TabICL and TabFlex—designed specifically to handle larger datasets—both underperformed naive TabPFN on genomic data, suggesting fundamental architectural limitations rather than just scalability issues. Classical methods with proper feature engineering achieved the best results (0.74 balanced accuracy).

### 4.3 Successful TabPFN Integration: Metadata-Based Subgrouping

Despite the failure of architectural adaptations, we discovered a successful approach to integrating TabPFN with genomic data by respecting rather than circumventing its design constraints. The key insight is to partition the large dataset into smaller, biologically meaningful subgroups based on metadata, then train separate TabPFN models on each subgroup.

#### 4.3.1 Organ-Specific TabPFN Models

We first partitioned samples by tissue type, training dedicated TabPFN models for each organ with sufficient sample size (*>* 300 samples). This approach naturally satisfies TabPFN’s sample size constraints while leveraging biological similarity within tissue types.

#### 4.3.2 Sex-Aware Stratification

For organs with large sample sizes (particularly blood with 20,000 samples), we further stratified by biological sex, training separate male and female TabPFN models. This additional stratification captures sex-specific gene expression patterns while maintaining sample sizes within TabPFN’s operational range.

Table 4 shows the substantial performance improvements achieved through this metadata-aware sub-grouping strategy.

**Table 4:** TabPFN Performance with Metadata-Based Subgrouping.

| Approach | Tissue | $R^2$ | MAE (years) |
| --- | --- | --- | --- |
| Naive TabPFN (blood only) | Blood | 0.657 | 8.9 |
| Naive TabPFN (colon only) | Colon | 0.7217 | 8.67 |
| Organ-Aware TabPFN | Multi-organ | 0.7188 | 8.25 |
| Sex-Aware TabPFN (blood only) | Blood | 0.705 | 8.175 |
| **Organ + Sex Aware TabPFN** | **Multi-organ** | **0.75** | **7.79** |

#### 4.3.3 Key Findings

The metadata-based subgrouping approach yields several important insights:

- **Organ-specific models outperform global models**: Colon-specific TabPFN (*R*^2^ = 0.7217) substantially outperforms naive TabPFN on the full dataset (*R*^2^ = 0.6074)
- **Sex stratification provides additional gains**: Sex-aware blood models achieve *R*^2^ = 0.705 compared to *R*^2^ = 0.657 for blood-only models
- **Combined strategy achieves best performance**: Organ + Sex Aware TabPFN reaches *R*^2^ = 0.75, representing a 23
- **Biological relevance matters**: Performance gains align with known biological differences between tissues and sexes in gene expression patterns

This approach demonstrates that TabPFN can be successfully applied to genomic data when its architectural constraints are respected through intelligent data partitioning rather than circumvented through complex adaptations.

## 5 Discussion

### 5.1 Failure of TabPFN Adaptations

Our results reveal fundamental limitations in adapting TabPFN for transcriptomics applications. Despite multiple sophisticated approaches—self-supervised pretraining, domain-specific architectures (BulkFormer), ensemble methods, and recent state-of-the-art TabPFN extensions (TabICL, TabFlex)—all TabPFN variants underperformed the naive baseline. This suggests that even with dimensionality reduction to 500 features, the resulting representations may lose critical biological signal. Moreover, RNA-seq datasets often have hundreds to thousands of samples, potentially insufficient for TabPFN’s in-context learning to be effective Also, TabPFN’s general-purpose design may be inherently unsuited for highly structured biological data

### 5.2 Self-Supervised Learning Limitations

The poor performance of self-supervised approaches (*R*^2^ = 0.494-0.5259) indicates that generic SSL objectives fail to capture relevant biological signal for age prediction. Masked gene modeling, contrastive learning, and VAE reconstruction may not align with the phenotypic prediction task, suggesting that task-specific feature learning is crucial for genomic applications.

### 5.3 Classical ML Superiority and TabPFN Viability

Classical methods achieve strong performance (3-layer MLP: *R*^2^ = 0.75), but our metadata-based subgrouping approach demonstrates that TabPFN can match this performance (Organ + Sex Aware TabPFN: *R*^2^ = 0.75, MAE = 7.79) when its constraints are respected. This suggests that the choice between TabPFN and classical methods for genomics depends on whether intelligent data partitioning is feasible and whether TabPFN’s in-context learning capabilities provide additional value for the specific application.

## 6 Conclusion and Future Work

We investigated multiple approaches for integrating TabPFN with high-dimensional RNA-seq analysis from our dataset of 57,873 samples with 10,000 genes each: self-supervised embedding learning, BulkFormer integration, ensemble methods, recent TabPFN extensions, and metadata-based subgrouping strategies.

Key findings include:

- **Architectural adaptation failures**: All complex TabPFN modifications (*R*^2^ = 0.494-0.627) underperformed naive TabPFN (*R*^2^ = 0.607), including recent state-of-the-art extensions TabICL and TabFlex
- **Successful constraint-respecting approach**: Metadata-based subgrouping by organ and sex achieves *R*^2^ = 0.75, matching classical ML performance while respecting TabPFN’s architectural limitations
- **Data strategy effectiveness**: Intelligent partitioning into biologically meaningful subgroups proves more effective than complex architectural modifications

Our results demonstrate that TabPFN’s 500-feature and 10,000-sample constraints cannot be effectively overcome through self-supervision or dimensionality reduction techniques. Instead, TabPFN excels when datasets are intelligently partitioned into smaller, biologically coherent subsets. Classical baselines demonstrate superior performance when utilizing large datasets and full feature spaces, while TabPFN provides fast and accurate predictions in small-sample, constrained-feature scenarios.

### Future Work

Several promising directions emerge from our analysis. First, developing biologically-informed feature selection methods that identify the most predictive gene subsets for specific phenotypes could better utilize TabPFN’s 500-feature capacity. Second, exploring hierarchical modeling approaches that combine multiple TabPFN models trained on different biological partitions may capture complex multi-organ interactions. Third, investigating domain-specific pre-training of TabPFN on large-scale genomic datasets could improve its inductive biases for biological applications. Finally, developing adaptive ensemble methods that automatically determine optimal data partitioning strategies based on dataset characteristics would enhance TabPFN’s practical deployment in computational biology.

Our work contributes to understanding optimal deployment scenarios for tabular foundation models in computational biology, demonstrating that respecting architectural constraints through intelligent data organization is more effective than attempting to circumvent them through complex adaptations.

## References

Yoav Benjamini and Yosef Hochberg. Controlling the false discovery rate: a practical and powerful approach to multiple testing. Journal of the Royal Statistical Society: Series B (Methodological), 57(1):289–300, 1995. doi: 10.1111/j.2517-6161.1995.tb02031.x.

Leo Breiman. Random forests. Machine Learning, 45(1):5–32, 2001. doi: 10.1023/A:1010933404324.

Corinna Cortes and Vladimir Vapnik. Support-vector networks. Machine Learning, 20(3):273–297, 1995. doi: 10.1007/BF00994018.

Noah Hollmann, Samuel Müller, Katharina Eggensperger, and Frank Hutter. Tabpfn: A transformer that solves small tabular classification problems in a second, 2023. URL https://arxiv.org/abs/2207.01848.

W. Evan Johnson, Cheng Li, and Ariel Rabinovic. Adjusting batch effects in microarray expression data using empirical bayes methods. Biostatistics, 8(1):118–127, 04 2006. ISSN 1465-4644. doi: 10.1093/biostatistics/kxj037. URL https://doi.org/10.1093/biostatistics/kxj037.

Ian Jolliffe. Principal Component Analysis, pages 1094–1096. Springer Berlin Heidelberg, Berlin, Heidelberg, 2011. ISBN 978-3-642-04898-2. doi: 10.1007/978-3-642-04898-2455. URL https://doi.org/10.1007/978-3-642-04898-2_455.

Boming Kang, Rui Fan, Meizheng Yi, Chunmei Cui, and Qinghua Cui. A large-scale foundation model for bulk transcriptomes. bioRxiv, 2025. doi: 10.1101/2025.06.11.659222. URL https://www.biorxiv.org/content/early/2025/06/17/2025.06.11.659222.

Diederik P. Kingma and Max Welling. Auto-encoding variational bayes. arXiv preprint 1312.6114, 2013.

Si-Yang Liu and Han-Jia Ye. Tabpfn unleashed: A scalable and effective solution to tabular classification problems, 2025. URL https://arxiv.org/abs/2502.02527.

Jingang Qu, David Holzmüller, Gaël Varoquaux, and Marine Le Morvan. Tabicl: A tabular foundation model for in-context learning on large data, 2025. URL https://arxiv.org/abs/2502.05564.

David E. Rumelhart, Geoffrey E. Hinton, and Ronald J. Williams. Learning representations by back-propagating errors. Nature, 323(6088):533–536, 1986. doi: 10.1038/323533a0.

Christina V. Theodoris, Ling Xiao, Anant Chopra, Mark D. Chaffin, Zeina R. Al Sayed, Michael C. Hill, Helene Mantineo, Elizabeth M. Brydon, Zexian Zeng, X. Shirley Liu, and Patrick T. Ellinor. Transfer learning enables predictions in network biology. Nature, 618(7965):616–624, 2023. doi: 10.1038/s41586-023-06139-9. URL https://doi.org/10.1038/s41586-023-06139-9.

Robert Tibshirani. Regression shrinkage and selection via the lasso. Journal of the Royal Statistical Society: Series B (Methodological), 58(1):267–288, 1996.

Laurens van der Maaten and Geoffrey Hinton. Visualizing data using t-sne. Journal of Machine Learning Research, 9(86):2579–2605, 2008. URL http://jmlr.org/papers/v9/vandermaaten08a.html.

Yuchen Zeng, Tuan Dinh, Wonjun Kang, and Andreas C Mueller. Tabflex: Scaling tabular learning to millions with linear attention, 2025. URL https://arxiv.org/abs/2506.05584.

